# Intracellular connections between basal bodies promote the coordinated behavior of motile cilia

**DOI:** 10.1101/2022.05.06.490816

**Authors:** Adam W. J. Soh, Louis G. Woodhams, Anthony D. Junker, Cassidy M. Enloe, Benjamin E. Noren, Adam Harned, Christopher J. Westlake, Kedar Narayan, John S. Oakey, Philip V. Bayly, Chad G. Pearson

## Abstract

Hydrodynamic flow produced by multi-ciliated cells is critical for fluid circulation and cell motility. Hundreds of cilia beat with metachronal synchrony for fluid flow. Cilia-driven fluid flow produces extracellular hydrodynamic forces that cause neighboring cilia to beat in a synchronized manner. However, hydrodynamic coupling between neighboring cilia is not the sole mechanism that drives cilia synchrony. Cilia are nucleated by basal bodies (BBs) that link to each other and to the cell’s cortex via BB-associated appendages. The intracellular BB and cortical network is hypothesized to synchronize ciliary beating by transmitting cilia coordination cues. The extent of intracellular ciliary connections and the nature of these stimuli remain unclear. Moreover, how BB connections influence the dynamics of individual cilia has not been established. We show by FIB-SEM imaging that cilia are coupled both longitudinally and laterally in the ciliate *Tetrahymena thermophila* by the underlying BB and cortical cytoskeletal network. To visualize the behavior of individual cilia in live, immobilized *Tetrahymena* cells, we developed <u>D</u>elivered Iron <u>P</u>article <u>U</u>biety <u>L</u>ive <u>L</u>ight-(DIPULL) microscopy. Quantitative and computer analyses of ciliary dynamics reveal that BB connections control ciliary waveform and coordinate ciliary beating. Loss of BB connections reduces cilia-dependent fluid flow forces.

**Summary:** Soh et al investigate whether intracellular connections between basal bodies control ciliary behavior in multi-ciliated cells. Using a *Tetrahymena* live cell immobilization technique to quantify ciliary dynamics, they show that inter-BB connections are required for effective ciliary waveform and coordinated ciliary beating that promotes fluid flow.

## Introduction

Hydrodynamic flow supports a myriad of biological functions, including the circulation of cerebrospinal fluid in the brain ventricles, mucosal clearance in the airway, transport of eggs in the oviduct, and cell motility in aquatic environments. In these specialized cell types, fields of motile cilia or multi-ciliary arrays decorate the cell surface. Motile cilia beat in a rhythmic and directed manner for unidirectional fluid flow.

Ciliary beating exerts motive forces on both the fluid environment and the cell body of multi-ciliary arrays. The direction and magnitude of cilia-driven forces are defined by several factors that include the ciliary waveform and ciliary beat frequency. The ciliary waveform refers to the propagation of bends along the cilium length during ciliary beating. According to the “sliding filament model”, ciliary bends are promoted by sliding between radially arranged and coupled doublet microtubules within the ciliary axoneme (Brokaw, 1972a; Brokaw, 1972b; Satir, 1968; Summers and Gibbons, 1971). Tight spatiotemporal regulation over this process is fundamental to the formation of unique ciliary waveforms (Naitoh and Sugino, 1984; Warner and Satir, 1974). During the power stroke, the cilium, which is akin to an oar of a boat, pushes fluid along the cell’s anterior-posterior axis while maintaining a relatively straight profile. This ensures that the magnitude of drag forces is maximized for effective propulsion of fluid along the cell body (Naitoh and Sugino, 1984). Concurrently, ciliary forces are also transmitted into the cell body. In motile multi-ciliated organisms, the opposing forces from fluid propulsion are utilized for cell motility. Once complete, the power stroke transitions into the recovery stroke whereby the cilium moves in the opposite direction. Notably, the cilium adopts a prominent bend while maintaining a low trajectory across the cell surface. This reduces drag forces on the cilium as it travels in the opposite direction. Then, the cycle repeats. The ciliary beat frequency, which reflects the speed of ciliary movement, also defines the magnitude of ciliary forces (Bottier et al., 2019; Naitoh and Sugino, 1984; Teff et al., 2008). Together, the direction and magnitude of cilia-driven forces depend on the ciliary waveform and ciliary beat frequency.

The coordination of ciliary beating is achieved by dynamic interactions between neighboring, oriented cilia (Elgeti and Gompper, 2013; Maestro et al., 2018). Hydrodynamic coupling synchronizes ciliary beating by interactions between cilia through their fluid environment (Elgeti and Gompper, 2013; Maestro et al., 2018; Tamm, 1984). Neighboring and oriented cilia undulate sequentially such that when one cilium beats, the fluid forces exerted upon its neighbor serve to coordinate their beating. Adjacent cilia beat with a temporal delay, or phase difference but remain in synchrony. The coordinated undulation of cilia appears as metachronal waves that propagate across the multi-ciliary array. Computer modeling suggests that hydrodynamic coupling is sufficient to promote coordinated ciliary behavior if cilia are near each other (Elgeti and Gompper, 2013; Riedel et al., 2005). Experimental and modeling data demonstrates that when separated by at least 50% of a cilium length, fluid-driven cilia coordination becomes less effective (Brumley et al., 2014). Consequently, this leads to inconsistent ciliary waveforms and inefficient fluid flow (Brumley et al., 2014; Elgeti and Gompper, 2013). While hydrodynamic coupling is integral to coordinated ciliary beating, it may not be the only mechanism for promoting ciliary synchrony. For example, metachronal waves traveling along the cell cortex continue to propagate across multi-ciliary arrays even when hydrodynamic flow is physically blocked (Narematsu et al., 2015). In addition, *Chlamydomonas* cells with cilia positioning defects fail to beat in coordination even when the cilia are positioned within hydrodynamic range (Hoops et al., 1984; Quaranta et al., 2015; Wan and Goldstein, 2016). This suggests that alternate mechanisms can coordinate ciliary movement along the cell cortex.

The cell cortex is thought to synchronize ciliary beating by transmitting cilia coordination cues through the cellular cortical cytoskeleton (Hyams and Borisy, 1975; Narematsu et al., 2015; Quaranta et al., 2015; Tamm, 1999). However, the nature of this cue has not been established. In the multi-flagellate *Koruga*, cortical contractions were proposed to promote “body waves”, which lead to the formation of metachronal waves across the cell cortex (Cleveland & Cleveland, 1966). However, subsequent studies suggested body waves did not arise from intracellular cortical contraction (Machemer 1974) and inhibition of ciliary beating in *Koruga* abolishes “body waves” (Tamm, 1999). This suggests that cilia-driven forces are transmitted into the cell cortex and this in turn induces “body waves”, opening the possibility that ciliary forces themselves provide intracellular coordination cues (Tamm, 1999). At the cell cortex, basal bodies (BB) nucleate and anchor cilia. BB-associated appendages mediate BB-BB and BB-cell cortex connections (Allen, 1967; Galati et al., 2014; Iftode et al., 1996; Jerka-Dziadosz et al., 1995; Junker et al., 2019; Soh et al., 2020; Vladar et al., 2012; Werner et al., 2011). In addition, BBs are embedded in the underlying cortical cytoskeleton that comprises microtubules, actin, and intermediate filaments (Gordon, 1982; Hard and Rieder, 1983; Lemullois, 1987; Mahuzier et al., 2018; Werner et al., 2011; Williams, 2004; Williams et al., 2006; Yasunaga et al., 2022). Hence, BBs, at the base of cilia, are physically interconnected by an underlying cytoskeletal matrix. Ciliary beating and linkages between BBs deform BBs and their associated accessory structures (Junker et al., 2022; Khanal et al., 2021; Vernon and Woolley, 2004; Warner and Satir, 1974). It is proposed that the extent of the connections between BBs and the cell cortex determines the magnitude of transmitted ciliary forces between BBs, which then shapes the ciliary waveform (Junker et al., 2022). Whether the transmission of ciliary forces through neighboring BBs serves as a mechanical cue for controlling ciliary waveforms and coordinating ciliary beating remains to be investigated directly.

While connections between BBs are required for coordinated ciliary dynamics in bi-flagellate and multi-ciliated organisms, the underlying mechanisms remain unclear. In the case of the bi-flagellated *Chlamydomonas*, the beat patterns of the two cilia or flagella are coordinated by striated fibers (SFs) that connect the BBs at the base of the flagella (Quaranta et al., 2015; Wan and Goldstein, 2016). The *Chlamydomonas* mutant, *vfl3*, disrupts SFs and displays abnormal beating patterns of the two flagella (Hoops et al., 1984; Quaranta et al., 2015; Wan and Goldstein, 2016). This would suggest that the coupling between BBs coordinates ciliary behavior. Notably, the loss of connections between the two BBs in *vfl3* mutants is also associated with BB disorientation, whereby BBs no longer dock at the cell cortex in a polarized manner (Hoops et al., 1984). Proper BB orientation in bi-flagellates and in multi-ciliary arrays are fundamental for polarized ciliary beating to drive fluid flow and cell motility (Soh and Pearson, 2021). It is plausible that disoriented BBs lead to ciliary beating defects, which consequently cause the loss of cilia coordination (Gilpin et al., 2017b; Quaranta et al., 2015; Wan and Goldstein, 2016). However, this has not yet been tested. In the case of vertebrate multi-ciliary arrays, the cortical actin network organizes BBs (Mahuzier et al., 2018; Werner et al., 2011). Loss of cortical actin causes cilia mispositioning and undocking from the cell cortex (Mahuzier et al., 2018; Werner et al., 2011). Moreover, synchronized ciliary beating is disrupted. Therefore, the cortical actin and BB network is not only required for cilia organization but is also necessary for normal ciliary dynamics. Finally, the metachronal behavior of multi-ciliary arrays in the ciliate, *Paramecium,* responds to physical manipulations to the cell cortex. Experimentally induced cyclical stretching along the *Paramecium* cell cortex changes their metachronal wave frequency (Narematsu et al., 2015). Together, this suggests that the cell cortex of multi-ciliary arrays also regulates ciliary coordination. However, the nature of the cortical elements, how forces are propagated between cilia, and the direct impact on ciliary beating is not understood.

## Results and Discussion

### *Tetrahymena* cilia are intracellularly coupled

To understand how the BB and cytoskeletal network synchronizes ciliary dynamics in multi-ciliary arrays, the interconnectedness of the underlying cortical architecture in *Tetrahymena* ciliary arrays was visualized. Individual *Tetrahymena* BBs establish connections with their BB neighbors and the cell cortex (Allen, 1967; Galati et al., 2014; Pitelka, 1961; Soh et al., 2020). However, the extent of these interactions across the entire cell surface has not been characterized. Using fluorescence microscopy and focused ion beam scanning electron microscopy (FIB-SEM) in *Tetrahymena* cells, we observed that each ciliary row is longitudinally connected along the entire cell length (Fig. 1). Consistent with prior work, each BB connection consists of an anteriorly oriented BB-associated striated fiber (SF) that is connected to the post-ciliary microtubule (pcMTs) bundle of the anterior BB (Figs. 1, S1A and Movie 1; (Allen, 1967; Galati et al., 2014; Junker et al., 2019; Soh et al., 2020)). SF-pcMT linkages are short (16.0±4.6 nm in length; (Soh et al., 2020)) and the resolution of FIB-SEM was not sufficient to visualize these structures. Therefore, we defined BB connections through SFs and pcMTs as those positioned within 1 SD of the average SF-pcMT linkage length (16.0±4.6 nm). BB connections also occur between ciliated and unciliated BBs, whereby the SFs of ciliated BBs interact with the pcMTs of anteriorly positioned unciliated BBs (Figs. 1B and S1B). In addition, ciliated BBs with long SFs can interact with the pcMTs of the anterior BB that are positioned two BBs away (Figs. 1B and S1B). Thus, BBs within a ciliary row are longitudinally connected (Fig. 1B). Between adjacent BB rows, the BB-associated transverse microtubules (tMTs) connect to the cell cortex in alignment with the adjacent ciliary row (Fig. S1A and Movie 1; (Junker et al., 2019)). The distal ends of tMTs point toward the right adjacent ciliary row (when viewed from above the cell) and either associate with or are near to (within 20 nm) the SFs and pcMTs of the adjacent ciliary row (Fig. S1A and Movie 1; (Junker et al., 2019; Soh et al., 2020; Williams, 2004)). Whether the tMT distal ends directly connect with these structures was not resolved in our images but suggests that such interactions exist. In summary, *Tetrahymena* cilia are longitudinally and laterally connected across the cell surface by an underlying BB and cortical cytoskeletal network (Figs. 1, S1A and Movie 1).

**Figure 1.**
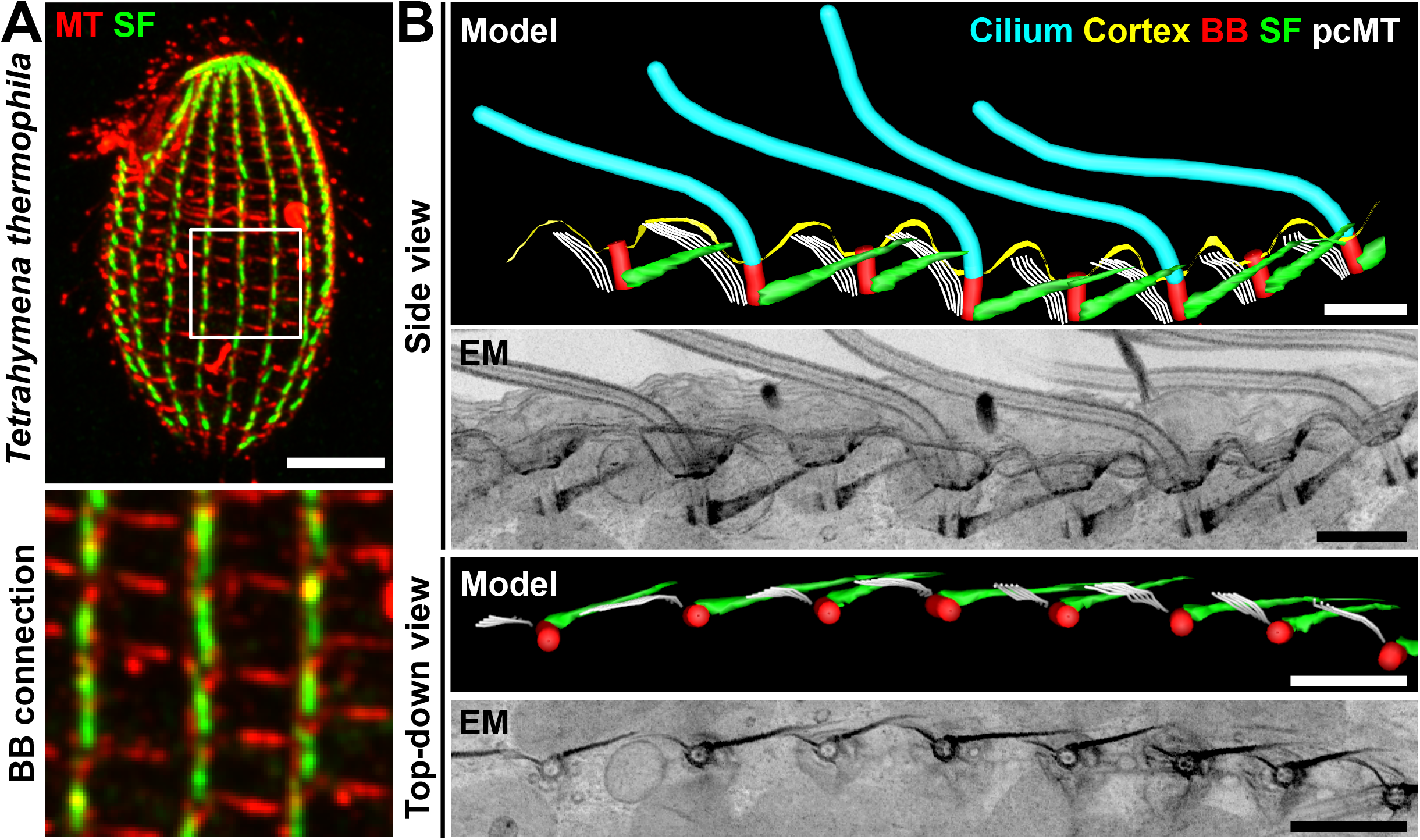
Cilia are intracellularly coupled by BBs. (A) Immunofluorescence image depicting a *Tetrahymena* cell. *Bottom panel*: Inset (white box) illustrating top-down views of the BB network. Microtubule (MT), red. SF, green. Scale bar, 10 µm. Inset width, 11.0 µm. (B) FIB-SEM (2D projection) and model images showing the intracellularly coupled ciliary array of *Tetrahymena* cell. Cilia, cyan; Cortex, yellow; BB, red; SF, green; Post-ciliary microtubule (pcMT), white. Scale bars, 1 µm.

The FIB-SEM analyses also showed that the long, proximal-distal axis of BBs is not uniformly perpendicular to the cell cortex. (Fig. S1C and D). This suggests that BBs rotate back and forth during ciliary beating in response to the mechanical forces from beating cilia. However, the limited sample size precluded our ability to determine whether BB angles directly correlate with cilia positions during their beat cycles (Fig. S1E). It is possible that BB rotations are caused by ciliary forces. At least two types of ciliary forces could cause BB movement. First, compressive and tensile forces from doublet microtubule sliding within the ciliary axoneme could impose forces directly on the BB (Junker et al., 2022; Khanal et al., 2021; Riedel-Kruse et al., 2007; Vernon and Woolley, 2004). Second and not exclusive of the first model, forces from neighboring ciliated BBs may be transmitted by intracellular BB connections. This is supported by the occurrence of unciliated BBs that are rotated relative to the cell cortex (Fig. S1C and D) and by our studies suggesting ciliary forces are transmitted to BBs by SFs (Junker et al., 2022). Further studies are required to uncover the source and magnitude of forces that promote BB rotation.

### Visualizing live *Tetrahymena* ciliary dynamics using DIPULL microscopy

To test whether connections between BBs are required for normal ciliary beating, we developed a strategy to visualize ciliary beating in live *Tetrahymena* cells. Rapid *Tetrahymena* movements during swimming obstructs the visualization and quantification of ciliary beating. We developed a method to immobilize live cells to visualize both extracellular fluid flow and ciliary dynamics. Prior live *Tetrahymena* cell immobilization strategies cause cytotoxicity or disrupt ciliary dynamics (Aufderheide, 1986; Kumano et al., 2012; Larsen and Satir, 1991; Spoon et al., 1977). To circumvent these issues, we developed <u>D</u>elivered Iron <u>P</u>article <u>U</u>biety <u>L</u>ive <u>L</u>ight (DIPULL) microscopy to trap live *Tetrahymena* cells (Fig. 2A). *Tetrahymena* cells were fed 1 - 2 µm iron particles that are engulfed by phagocytosis (Figs. 2A (Step 1) and S2A; (Rifkin and Ballentine, 1976)). Cells were then trapped and immobilized within PDMS microfluidic chambers using magnetism and then visualized using an inverted microscope (Figs. 2A (Step 2) and S2J). DIPULL does not compromise ciliary dynamics, cell morphology, or cell division (Fig. S2A - H). This immobilization strategy traps live *Tetrahymena* cells allowing for quantitative visualization of ciliary dynamics.

**Figure 2.**
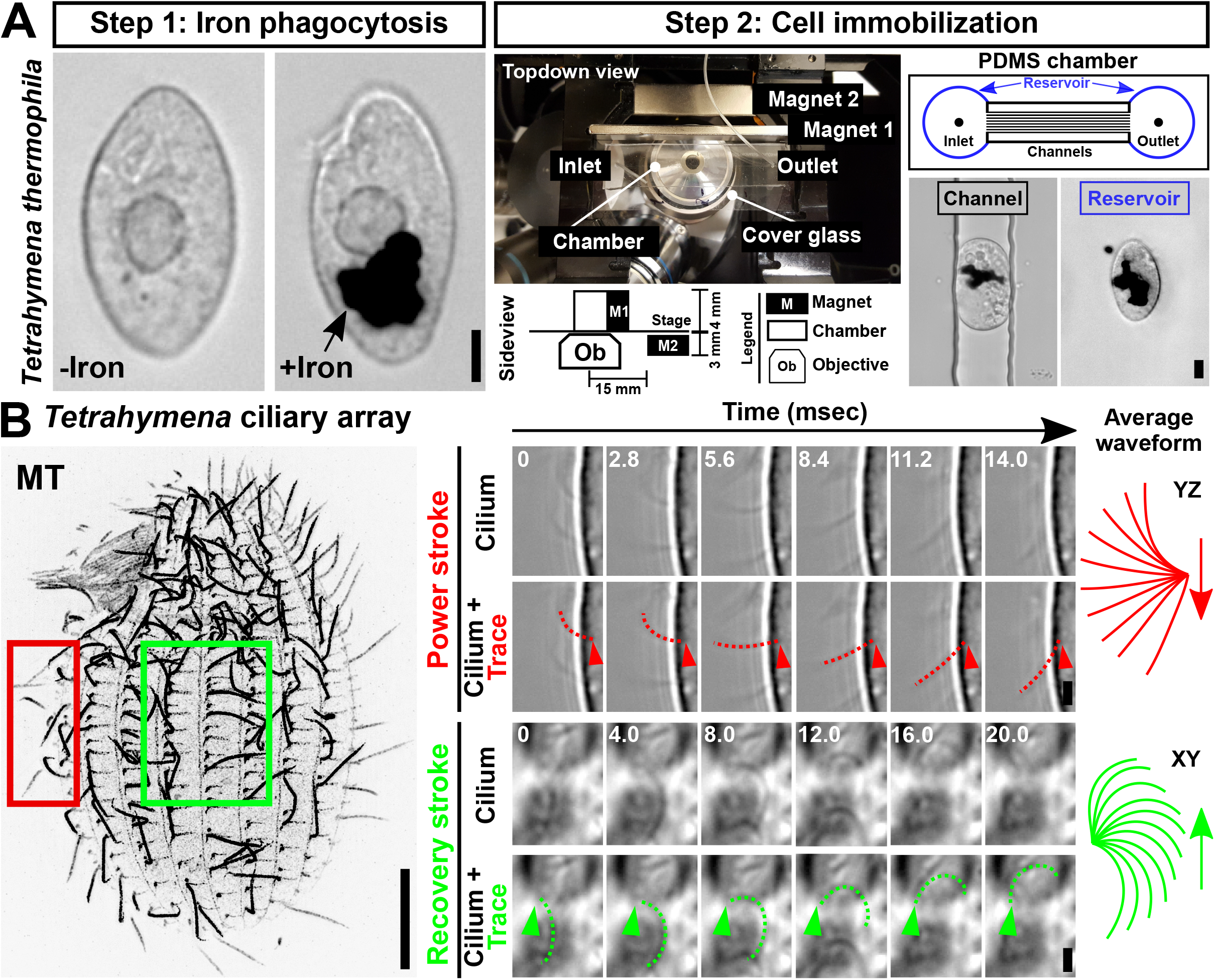
*Tetrahymena* live cell immobilization technique. (A) <u>D</u>elivered Iron <u>P</u>article <u>U</u>biety <u>L</u>ive <u>L</u>ight (DIPULL) - microscopy setup. *Step 1*: *Tetrahymena* cells are fed iron particles. Cell images pre- and post-iron engulfment. *Step 2*: Cells are introduced into a microfluidic chamber and immobilized via a constant external magnetic field. To track intracellular dynamics, imaging was performed on cells that were trapped within channels (black outline). The visualization of extracellular dynamics was performed on cells that are trapped within the chamber reservoir (blue outline). Scale bars, 10 µm. (B) Visualization of ciliary dynamics via DIPULL-immobilized live *Tetrahymena* cells. *Left panel*: *Tetrahymena* ciliary array. Bar, 10 µm. *Right panel*: Timelapse images of power and recovery strokes. Time intervals (msec) are indicated. Dotted lines mark manual cilia traces. Average power stroke, red. Average recovery stroke, green. Scale bar, 2 µm.

### BB connections and orientation are required for efficient fluid flow

Using DIPULL, we tested whether BB disconnection and disorientation affects cilia-dependent fluid flow. To investigate whether cilia-driven fluid flow is disrupted when BB connections and orientation are lost, fluid flow was quantified in WT and *disA-1* cells (Fig. 3A). *disA-1* mutants swim slower and possess short SFs where 82% of BBs are disconnected from their neighbors (Fig. 3A; (Galati et al., 2014; Nabi et al., 2019; Soh et al., 2020)). Cells were immobilized in the DIPULL chamber reservoir to avoid boundary effects from the chamber walls (Fig. 2A). Fluid flow was visualized by tracking the movement of 0.5 µm fluorescent beads that were added to the media (Fig. 3B and C). The movements of beads were visualized by projecting images from a time-lapse of 2.5 sec (100 frames; Movie 2). Sustained fluid flow traveling from the cell’s anterior (A) pole to the posterior (P) pole was observed in WT cells (Fig. 3B (left) and Movie 2 (left)). Flow was faster at the cell anterior pole compared to the posterior pole (Fig. 3B (right); WT mean fluid flow velocity: anterior pole: 26.0±8.6 µm/ sec; posterior pole: 14.0±9.9 µm/ sec; Mann-Whitney test; P value < 0.001). *disA-1* cells with disconnected and disoriented BBs exhibit shorter fluorescent bead movements, indicating that fluid flow is slower (Fig. 3C (left); Movie 2 (right)). Like WT cells, *disA-1* cells have faster fluid flow at the cell anterior compared to the posterior (Fig. 3C (right); *disA-1* mean fluid flow velocity: anterior pole: 11.0±9.0 µm/ sec; posterior pole: 5.6±5.9 µm/ sec; P value = 0.002). Consistent with the reduced swimming speed of *disA-1* mutants, the fluid flow velocities are approximately 50% of WT cells (Fig. 3B and C; Mann-Whitney test; P value < 0.001; (Galati et al., 2014)). This suggests that BB connections may be important for sustained fluid flow (Fig. 3B and C). However, in addition to being disconnected, 63% of BBs in *disA-1* cells are disoriented, making it impossible to distinguish whether the observed fluid flow reduction results from BB disconnections and / or BB disorientation.

**Figure 3.**
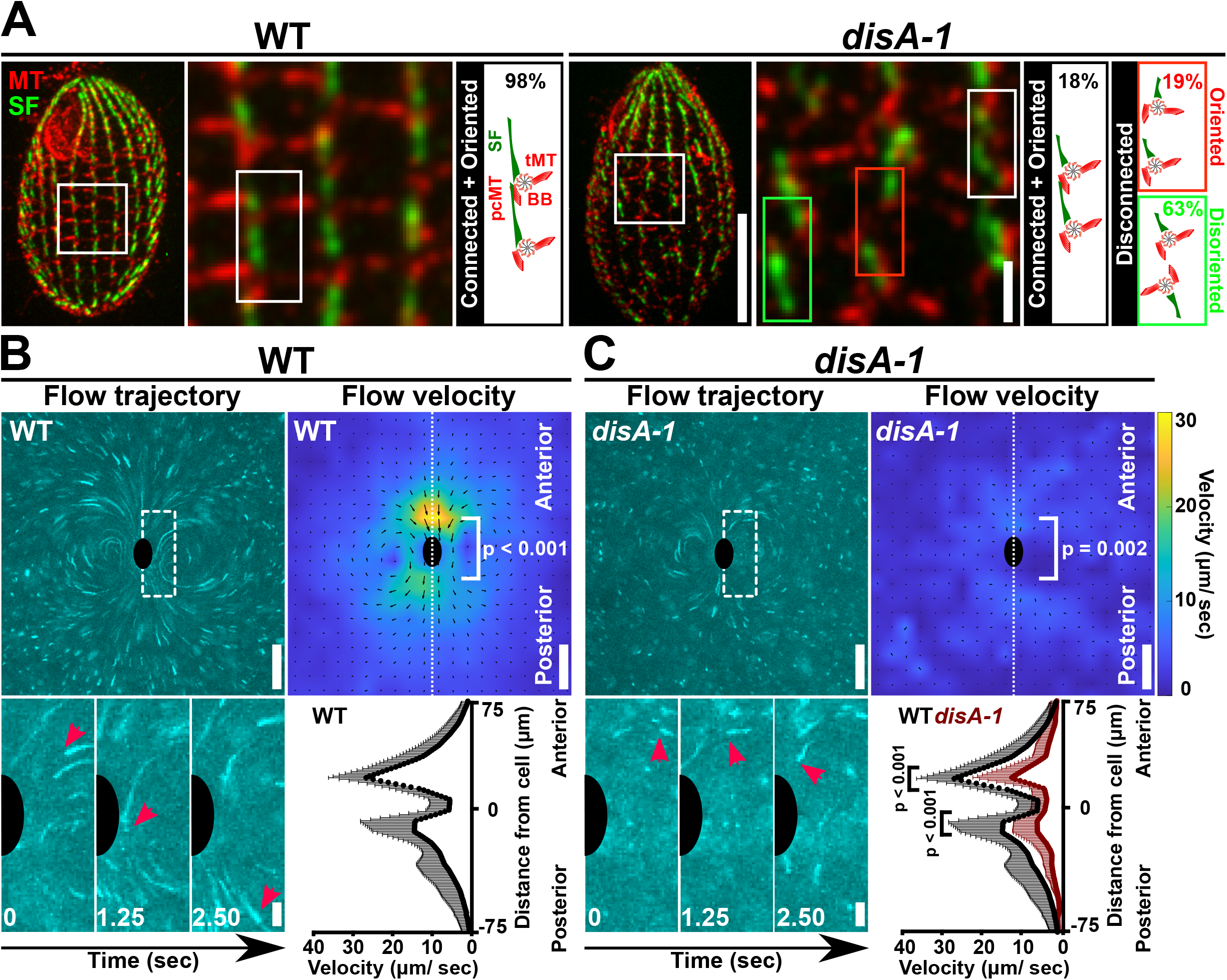
BB connections and orientation are required for cilia-dependent fluid flow. (A) Fluorescence images depicting the cortical organization of WT and *disA-1* cells at 25°C. Most *disA-1* cells display BB disconnection (82%). Disconnected *disA-1* bodies are either oriented (19% of total) or disoriented (63% of total). Microtubule (MT), red. SF, green. Scale bars, 10 µm and 2 µm. (B) BB connections are required for normal fluid flow movements and velocity at 25°C. Fluid flow is assessed by tracking fluorescent beads in media. Fluid flow trajectories are depicted as time projected images over 2.5 sec (100 frames). Inset (white box with dashed line) illustrates the movement of fluorescent bead (red arrowheads) relative to the cell (black half oval). Fluid flow velocity is represented as heatmaps. Cooler colors indicate slower fluid flow while warmer colors indicate faster fluid flow. Predicted cell position is marked by a black oval. (C) *disA-1* disrupts fluid flow velocity at 25°C. Inset (white box with dashed line) illustrates the movement of fluorescent bead (red arrowheads) relative to the cell (black half oval). WT (black line): n = 9 cells. *disA-1* (red line): n = 13 cells. Mann-Whitney test. Mean±SD. Scale bars, 50 µm (full field of view) and 10 µm (inset).

### Loss of BB connections reduces ciliary forces

To test whether BB connections specifically impact ciliary forces, we focused on cilia with BBs that are disconnected but still oriented. Oriented cilia were identified as those in which the power stroke occurs along the cell’s anterior-posterior axis and remain in-focus during the entire power stroke before transiting into the recovery stroke where they exit the imaging focal plane (Fig. 2B (right) and Movie 3 (WT: left; *disA-1* oriented cilia: middle; *disA-1* disoriented cilia: right)). We also imposed the requirement that the anterior and posterior neighboring cilia are similarly oriented (Movie 3). Of the BBs acceptable for analysis, we predict that 51% will be disconnected from neighboring BBs (Fig 3A (right)). Ciliary parameters that affect fluid flow, are defined at minimum by ciliary length, ciliary beat frequency or speed, ciliary sweep trajectory path, and ciliary curvature. Ciliary length is not changed in *disA-1* mutants (Fig. S2I; WT: 5.0±0.7 µm; *disA-1*: 5.2±0.6 µm; Student’s t-test; P = 0.35), and we therefore focused on the remaining three parameters to determine if they were impacted when BBs are disconnected.

#### Loss of BB connections slows the power stroke but does not impact the overall ciliary beat frequency

To test whether the disruption of BB connections specifically reduces ciliary beat frequency and ciliary speed, ciliary beat frequency was quantified using kymograph analysis of oriented cilia viewed from the side (Fig. 4A). The average ciliary beat frequency of WT cells at 25°C is 24.0±5.7 Hz (Fig. 4B and 4C). Faster ciliary beat frequencies were previously reported for *Tetrahymena* cells (Stoddard et al., 2018) and we suggest this is attributed to differences in strain background, the position of the cilia along the cell cortex, and reduced temperature. The average ciliary beat frequency of both oriented and disoriented *disA-1* cilia is comparable to WT cilia (Figs. 4C and S3C; ciliary beat frequency; WT: 24.0±5.7 Hz; oriented *disA-1*: 24.0±7.4 Hz; disoriented *disA-1*: 23.0±7.8 Hz; Mann-Whitney test; P = 0.55 and P = 0.94, respectively). The distribution of ciliary beat frequency in the disoriented cilia appears bimodal, whereby a subpopulation of cilia has reduced ciliary beat frequencies, but the overall mean ciliary beat frequency is like that of oriented cilia (Fig. S3C). Ciliary beat frequency was next imaged from above the cell to track the ciliary power stroke trajectory (Fig. S3A). Consistent with results using side views, top views of oriented *disA-1* cilia had comparable ciliary beat frequencies to WT cilia (Fig. S3B; WT: 24.0±4.8 Hz; *disA-1*: 25.0±7.6 Hz; Mann-Whitney test; P = 0.51). These results suggest that BB connections and orientation are not required for the overall ciliary beat frequency (Figs. 4C and S3C). While *Chlamydomonas vlf3* mutants with disconnected BBs exhibit variable and reduced ciliary beat frequency (Wan and Goldstein, 2016), *Tetrahymena disA-1* mutants appear to have relatively normal ciliary beat frequencies (Figs 4C and S4B, C).

**Figure 4.**
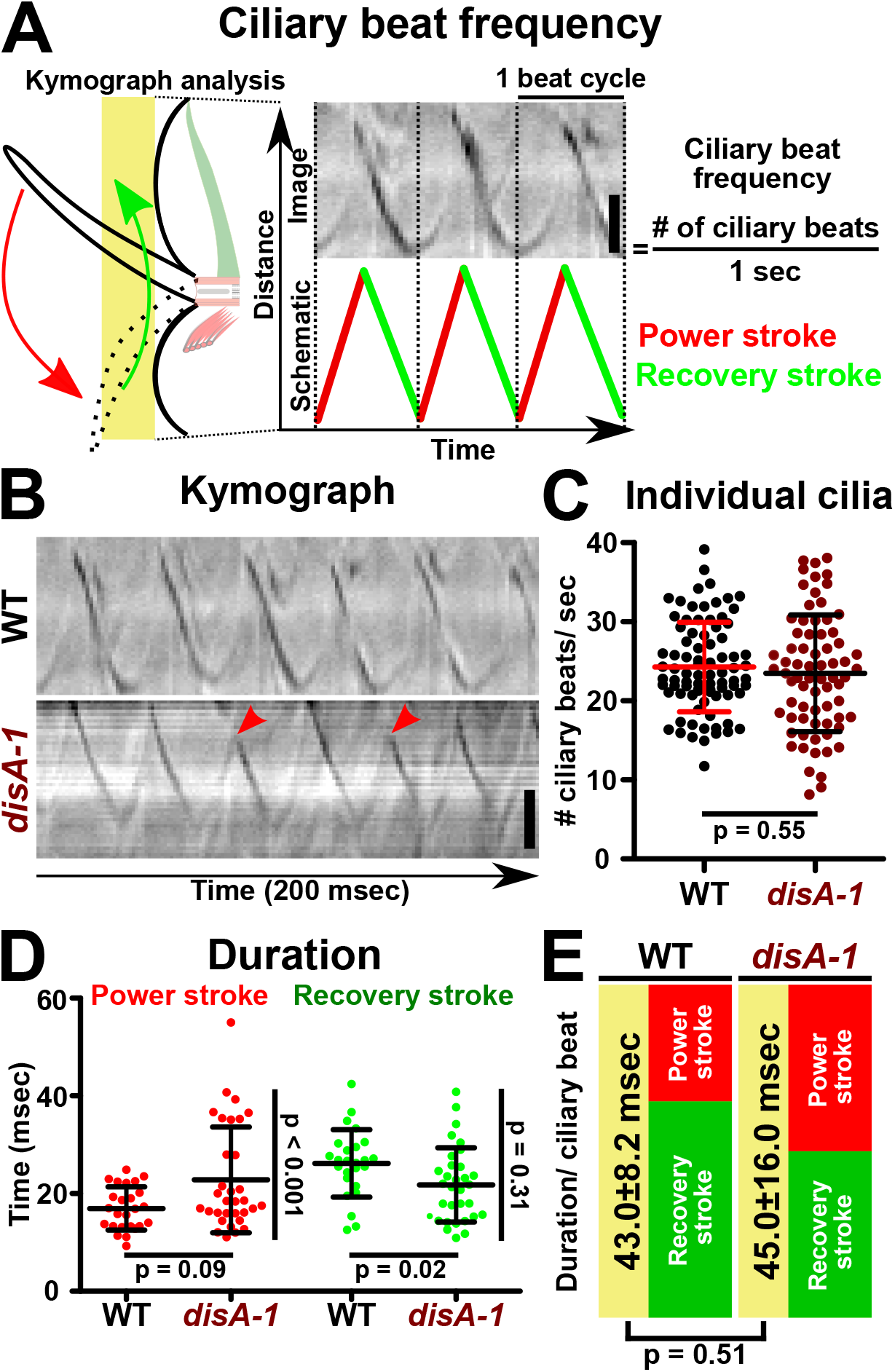
Loss of BB connections does not inhibit ciliary beat frequency. (A) Schematic image depicting ciliary beat frequency quantification using kymograph analysis. X-axis represents time (total duration: 105 msec). Y-axis represents distance. Scale bar, 2 µm. (B) Kymographs of WT and *disA-1* cilia. *disA-1* cilia display variable ciliary trajectories between ciliary beat cycles (red arrowheads). Scale bar, 2 µm. (C) Quantification of WT and *disA-1* ciliary beat frequency of individual cilia. Three cilia per cell were sampled. *disA-1* cilia exhibit comparable ciliary beat frequencies to WT cilia. WT: n = 28 cells (84 cilia). *disA-1*: n = 38 cells (72 oriented cilia; 41 disoriented cilia). Mann-Whitney test. Mean±SD. (D) Duration of power and recovery strokes of WT and *disA-1* cilia. WT: n = 11 cells (25 cilia). *disA-1*: n = 13 cells (32 cilia). Mann-Whitney test. F-test for variance comparison. Mean±SD. (E) Schematic illustrating comparable average duration of ciliary beat cycle for WT and *disA-1* cilia. *disA-1* cilia beat at comparable ciliary beat frequency as WT cilia with a longer power stroke duration and a shorter recovery stroke duration.

WT and *disA-1* cilia tips were imaged from above the cell to quantify the relative durations of the power and recovery strokes. The power stroke duration of WT and *disA-1* cilia is 17.0±4.4 msec and 23.0±11.0 msec, respectively (Fig. 4D; Mann-Whitney test; P value = 0.09). The *disA-1* power stroke duration was more variable with a subpopulation of cilia exhibiting a longer duration power stroke (Fig. 4D; F-test for variance comparison; P < 0.001). This variability may reflect the mixture of connected and disconnected BBs present in the analysis (Fig. 3A), as we are not able to determine whether BBs are connected in our live imaging approach. In summary, the average *disA-1* power stroke duration is longer than WT. The recovery stroke of *disA-1* cilia is 4.0 msec shorter than WT (Fig. 4D; WT: 26.0±6.9 msec; *disA-1*: 22.0±7.6 msec; P value = 0.02). Thus, the average total duration of each ciliary beat cycle is comparable between WT and *disA-1* cilia (Fig. 4E; average total duration per ciliary beat cycle; WT: 43.0±8.2 msec (23.2 Hz); *disA-1*: 45.0±16.0 msec (22.2 Hz); Mann-Whitney test; P value = 0.51) and is consistent with the ciliary beat frequency.

The above measurements are based on tracking the ciliary tip thereby making quantification of the ciliary waveform through the ciliary beat cycle impossible. To quantify the ciliary generated forces, both ciliary sweep angle trajectory and cilia curvature must be quantified (see below), which requires the entire cilium to be within the imaging focal plane. The relative start and end positions of the power stroke were defined as when cilia first appear and exit the side view imaging focal plane, respectively (Fig. 2B; red box). The same method was applied to define the average start and stop positions of the recovery stroke when cilia were imaged from the top of the cell (Fig. 2B; green box). Importantly, this method only captures a portion of the complete power and recovery strokes when cilia are in focus. The WT power stroke duration was 18.0±9.3 msec while that of *disA-1* cilia was 11.0±1.8 msec (Mann-Whitney test; P value = 0.018). While the WT power stroke duration is similar to the above analyses, the *disA-1* power stroke is much shorter and likely represents a shorter duration in the imaging plane. The WT and *disA-1* recovery stroke durations were similar to each other but significantly reduced compared to the above analyses (WT = 10.0±4.3 msec and *disA-1* = 10.0±3.1 msec; Mann-Whitney test; P value = 0.73). Like the power stroke, this shorter measured duration likely results from the recovery stroke moving out of the imaging focal plane. These limited segments of the ciliary beat cycle were next used to quantify the ciliary waveforms and predicted force outputs.

#### BB connections support normal ciliary waveforms

The sweep trajectory of *disA-1* cilia is not always consistent between beat strokes (Fig. 4B; red arrowheads), suggesting that the ciliary waveform is disrupted when BBs are disconnected. Both the power and recovery strokes were imaged and quantified using a semi-automated image analysis regime (Figs. 2B, 5A and Movies 3 and 4 (Bottier et al., 2019)). Generally, the ciliary trajectories for WT power and recovery strokes are comparable to prior qualitative studies (Fig. 5A and F; (Fabritius et al., 2021; Joachimiak et al., 2021; Urbanska et al., 2018; Wood et al., 2007)).

**Figure 5.**
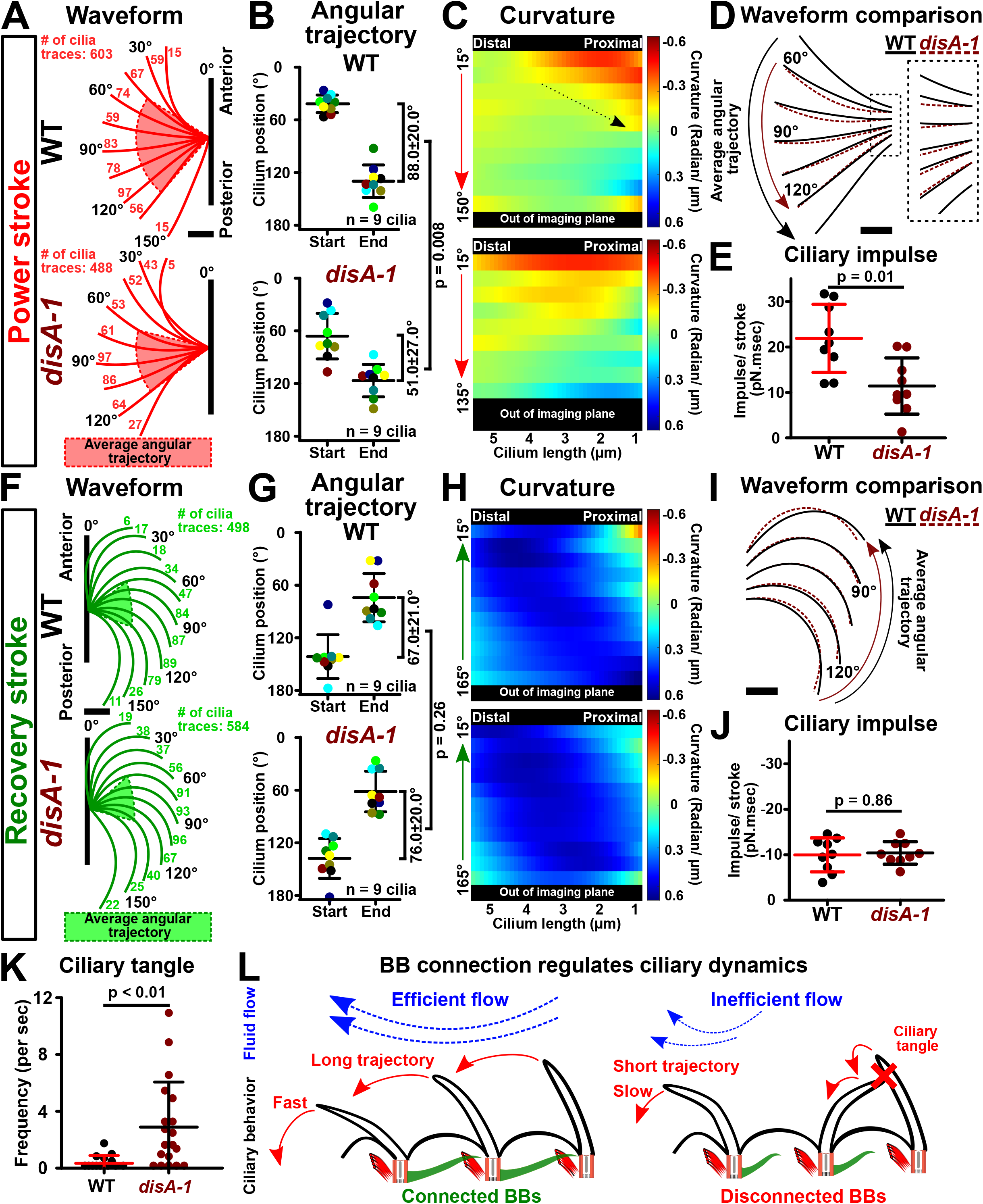
BB connections support ciliary waveform and coordination. (A) *Tetrahymena* power stroke waveform. *Top panel*: Average WT power stroke waveform. *Bottom panel*: Average *disA-1* power stroke waveform. Red highlight indicates the average angular trajectory (n = 9 cilia). Cilium position is defined by the angle from the cilium distal end (4.5 µm up the cilium base) relative to the cell’s anterior-posterior axis. Angles are categorized into 15° bins. Each angular bin is at least 1% of all ciliary traces per condition. Number of cilia traces for each bin is indicated. (B) Angular trajectories of WT and *disA-1* cilia. *disA-1* cilia undergo shorter trajectories than WT cilia during the power stroke. Each cilium in the analysis is color coded. (C) Curvature heatmaps of WT and *disA-1* cilia through the power stroke (blue: cilium bends towards the cell anterior pole; red: cilium bends away from the cell anterior pole; green: straight cilium). Y-axis marks the cilium position through the power stroke. Dotted arrow indicates the change in WT cilium curvature at the start to middle of the power stroke. WT power stroke: n = 603 cilia traces (9 cilia, 6 ciliary beat cycles each, 9 cells). *disA-1* power stroke: n = 488 cilia traces (9 cilia, 9 cells). (D) Superimposed WT (black lines) and *disA-1* (brown dashed lines) average power stroke waveform. Curvature differences are depicted only for stages of the power stroke that fall within the average angular trajectories of WT and *disA-1* cilia. (E) Ciliary power stroke impulses were estimated using resistive force theory. WT cilia exert greater impulse (area under the force-time curve) than *disA-1* cilia along the anterior-posterior axis per power stroke (n = 9 cilia). (F) *Tetrahymena* recovery stroke waveform. *Top panel*: Average WT recovery stroke waveform. *Bottom panel*: Average *disA-1* recovery stroke waveform. Green highlight indicates the average angular trajectory (n = 9 cilia). Cilium position is defined by the angle from the cilium distal end (4.5 µm up the cilium base) relative to the cilium’s power stroke axis. Angles are categorized into 15° bins. Each angular bin is at least 1% of all ciliary traces per condition. Number of cilia traces for each bin is indicated. (G) Angular trajectories of WT and *disA-1* cilia. Each cilium in the analysis is color coded. *disA-1* cilia undergo comparable trajectories as WT cilia during the recovery stroke. (H) Curvature heatmaps of WT and *disA-1* cilia through the recovery stroke (blue: cilium bends towards the cell anterior pole; red: cilium bends away from the cell anterior pole; green: straight cilium). Y-axis marks the cilium position through the recovery stroke. WT recovery stroke: n = 498 cilia traces (9 cilia, 9 cells). *disA-1* recovery stroke: n = 584 cilia traces (9 cilia, 6 ciliary beat cycles each, 9 cells). (I) Superimposed WT (black lines) and *disA-1* (brown dashed lines) average recovery stroke waveform. Curvature differences are depicted only for stages of the recovery stroke that fall within the average angular trajectories of WT and *disA-1* cilia. (J) Ciliary recovery stroke impulses were estimated using resistive force theory. WT and *disA-1* cilia exert comparable impulse (area under the force-time curve along the anterior-posterior axis per recovery stroke (n = 9 cilia). (K) Frequency of ciliary tangles. WT: n = 11 cells (14 cilia pair); *disA-1*: n = 13 cells (19 cilia pairs). (L) Schematic illustrates model that BB connections promote fast and long power stroke trajectories for coordinated ciliary beating and effective fluid flow propulsion. Scale bars, 1 µm.

Quantitative comparison between the ciliary waveforms of oriented WT and *disA-1* cilia suggest that cilia display a shorter angular sweep trajectory along the power stroke compared to WT cilia (Figs. 5A - B and S3I; WT cilia: 88.0±20.0°; *disA-1* cilia: 51.0±27.0°; Mann-Whitney test; P = 0.008). These average representative waveforms are generated from many individual ciliary traces that are standardized by using the average start and stop positions of all analyzed cilia (Fig. 5A (red highlights) and B; n = 6 beat cycles of 9 cilia total). *disA-1* cilia initiate the power stroke at 66.0±26.0° compared to WT cilia at 42.0±10°; and then transition into the recovery stroke at 120.0±18.0° compared to 130.0±19.0° for WT cilia (Fig. 5F (green highlights) and G). Subtle increases in variance were observed in *disA-1* cilia and may be explained by the subpopulation (49%) of oriented cilia with connected BBs that we expect to behave like WT cilia. The *disA-1* power stoke is shorter, suggesting that BB connections promote a long ciliary power stroke trajectory (Figs. 4D, 5A - B and S3I).

Changes to ciliary curvature alter the efficiency of fluid flow as straight cilia promote greater fluid propulsion than bent cilia (Brokaw, 1972a; Gray and Hancock, 1955; Naitoh and Sugino, 1984). We assessed the impact of BB connections on cilia curvature. A curvature value of zero indicates a straight cilium, a negative curvature value indicates a concave cilium profile that bends away from the cell anterior, and a positive curvature value indicates a convex cilium profile that bends towards the cell anterior. The greatest differences between WT and *disA-1* ciliary curvature occur primarily at the start to the middle of the power stroke (Figs. 5D and S3D). WT cilia are approximately 12-fold more bent than *disA-1* cilia at the proximal end of the cilium (Fig. 5D (inset); WT: −0.17±0.05 radians/ µm; *disA-1*: 0.01±0.11 radians/ µm; Mann-Whitney test; P < 0.0001). Importantly, the most proximal region of cilia (1.5 µm) could not be detected in our imaging. There may be unobservable differences between WT and *disA-*1 in this proximal region. At the medial region of the cilium, *disA-1* are 2-fold more bent than WT cilia (Fig. 5D; WT: −0.08±0.02 radians/ µm; *disA-1*: −0.14±0.04 radians/ µm; Mann-Whitney test; P < 0.0001). Together, this suggests that BB connections regulate ciliary curvature during the power stroke (Figs. 5A − D and S3D).

The recovery stroke returns cilia back to the start of the power stroke. Unlike the power stroke, BB connections do not appear to impact the angular trajectory of the recovery stroke (Fig. 5G and S3J; WT cilia: 67.0±21.0°; *disA-1* cilia: 76.0±20.0°; Mann-Whitney test. P = 0.26). We were unable to visualize the transition from the power to the recovery stroke and vice-versa because cilia move out of the focal plane during the 3-dimensional ciliary beat stroke. Seemingly contradictory, the recovery stroke trajectory of WT and *disA-1* cilia appear to be similar despite the shorter average sweep trajectory of the *disA-1* power stroke (Fig. 5B and G). One explanation is that the recovery stroke imaging plane of WT and *disA-1* cilia is different whereby WT cilia move closer to the cell cortex. This would result in comparable recovery stroke trajectory angles between WT and *disA-1* cilia, but the total distance traveled would be reduced for *disA-1* cilia. Alternatively, *disA-1* cilia may change their trajectory during transitions between power and recovery strokes outside the imaging focal plane. We next measured the difference in curvature in *disA-1* cilia and found they are slightly more bent during the recovery stroke than WT (Fig. 5I; WT: 0.46±0.11 radians/ µm; *disA-1*: 0.51±0.10 radians/ µm; Mann-Whitney test; P < 0.0001). Thus, our results suggest that BB connections are not required for normal recovery stroke angular sweep trajectories but do increase ciliary curvature (Figs. 5I, S3E and J).

#### Ciliary changes resulting from BB disconnections reduce ciliary-driven force

To test whether the shorter angular sweep trajectory and abnormal curvature exhibited by *disA-1* cilia during the power stroke reduce ciliary effectiveness, the magnitude of force along the anterior-posterior axis was estimated using resistive force theory (RFT). RFT uses resistive force coefficients to determine the distributed viscous drag applied to the cilium by the surrounding fluid based on the orientation and velocity of the cilium as a function of arc length and time (Gray and Hancock, 1955). Overall force produced by a cilium during its beat stroke will be affected by the cilium length, angular sweep rate, beat shape, and viscosity of the medium. WT cilia exerted an average force of 1.41±0.60 pN during the observed power stroke (Fig. S3F and Table 1; maximum power stroke force by WT cilia: 2.70±0.88 pN). The average force exerted during the observed power stroke by *disA-1* cilia is reduced by 18% (Fig. S3F and Table 1; *disA-1*; average power stroke force: 1.16±0.79 pN; maximum power stroke force: 2.11±1.19 pN). As shown in Fig. 5D, differences in curvature of WT and *disA-1* cilia are concentrated at the beginning of the power stroke, whereas differences in force output are greater toward the middle and end of the power stroke. This suggests that the reduced force of *disA-1* cilia is not due to the change in curvature but rather to the reduced angular sweep rate (Table 1; WT: 6.62±2.72°/ msec; *disA-1*: 5.20±2.88°/ msec). The magnitude of force production in the direction opposing cell motion during the recovery stroke was similar between WT and *disA-1* cilia (Fig. S3G and Table 1; average recovery stroke force; WT: −1.0±0.3 pN; *disA-1*: −1.1±0.4 pN; Mann-Whitney test; P value = 0.68).

**Table 1.** Tabulation of results of ciliary force estimation. Ciliary forces were estimated from manual traces using previously described resistive force coefficients (*C_N_=1.53µ*, *C_T_=0.64µ*; (Bayly et al., 2011)). Viscosity of water, *µ*, applied in analysis was 0.889cP at 25°C. All average values for ciliary quantities were calculated by averaging values within the mean sweep angle range for each traced ciliary beat stroke (54 total beat strokes each in 4 data sets), then calculating the average and standard deviation of the average quantities. Maximum values were calculated by taking the maximum value for each stroke within the mean sweep angle range, then calculating the average and standard deviation for the maximum quantities. All force values integrate forces in the anterior-posterior axis direction over the length of the cilium trace. Sweep rates are calculated by numerically differentiating the angle from the base of the cilium trace to 4.5 µm up the cilium with respect to time. Calculation of basal moments and power is described in *Materials and Methods – Ciliary Force Estimation*. WT and *disA-1* swim speeds (Galati et al., 2014) were used to estimate viscous drag forces on the *Tetrahymena* cell body using previously described methods (Bayly et al., 2011; Chwang and Wu, 1975).

| Parameter | WT | *disA-1* | P value |
| --- | --- | --- | --- |
| Average power stroke force [pN] | 1.41 ± 0.60 | 1.16 ± 0.79 | 0.07 |
| Maximum power stroke force [pN] | 2.70 ± 0.88 | 2.11 ± 1.19 | 0.001 |
| Average recovery stroke force [pN] | -1.05 ± 0.34 | -1.13 ± 0.40 | 0.73 |
| Maximum recovery stroke force [pN] | -1.51 ± 0.51 | -1.68 ± 0.56 | 0.15 |
| Average power stroke sweep rate [degree/ msec] | 6.62 ± 2.72 | 5.20 ± 2.88 | 0.04 |
| Maximum power stroke sweep rate [degree/ msec] | 10.69 ± 4.18 | 7.74 ± 4.00 | 0.002 |
| Average recovery stroke sweep rate [degree/ msec] | 5.83 ± 1.94 | 6.37 ± 2.28 | 0.31 |
| Maximum recovery stroke sweep rate [degree/ msec] | 7.60 ± 2.02 | 9.13 ± 3.11 | 0.008 |
| Average power stroke basal moment [pN·µm] | 2.86 ± 0.87 | 1.93 ± 1.44 | 0.00003 |
| Maximum power stroke basal moment [pN·µm] | 10.58 ± 3.16 | 8.37 ± 5.05 | 0.0005 |
| Average recovery stroke basal moment [pN·µm] | 1.13 ± 0.59 | 1.36 ± 0.72 | 0.16 |
| Maximum recovery stroke basal moment [pN·µm] | 4.90 ± 1.98 | 5.10 ± 1.93 | 0.44 |
| Average power stroke power [aW] | 870 ± 600 | 690 ± 640 | 0.02 |
| Average recovery stroke power [aW] | 420 ± 220 | 500 ± 310 | 0.27 |
| Swimming Speed [µm/ sec] | 272 | 123 | - |
| Body force / cilium [pN] | -0.26 | -0.12 | - |

Not only do *disA-1* cilia produce lower power stroke forces on average, but they also appear to spend a shorter duration of time producing that force due to the shorter angular sweep trajectory in our segmented analysis using only cilia in focus (Fig. 5A and B). This leads to a lower *impulse*, the measure of the area under the force-time curve. The mean axial impulse of WT cilia was 21.9±7.5 pN·ms, and that of *disA-1* cilia was 11.4±6.2 pN·ms (Fig. 5E, Mann-Whitney test; P = 0.01). During the recovery stroke, WT cilia produce an impulse of −10.0±3.7 pN·ms and *disA-1* cilia produce an impulse of −10.4±2.5 pN·ms (Fig. 5J, Mann-Whitney test; P = 0.86). Though the power stroke of *disA-1* cilia is significantly less productive, the recovery stroke produces a similar amount of drag impulse. The average measured impulses sum to 11.9±8.4 pN·ms per beat for WT cilia, and 1.0±6.7 pN·ms per beat for *disA-1* cilia, providing an estimate of the overall effectiveness of each beat. It should be noted that it was not possible to measure power and recovery strokes on the same cilia, and that the standard deviations on the power and recovery stroke impulses for *disA-1* cilia are larger than the difference in their magnitudes. For WT cilia, multiplying the net average impulse of 11.9 pN·ms per beat by the average frequency of 24 Hz gives an average total force per cilium of 0.29 pN, which compares well to the body drag force per cilium of −0.26 pN (Table 1). In summary, *disA-1* cilia are less effective, in part due to their lower average force output during their beat stroke, but more significantly due to the shorter time duration spent sweeping a shorter trajectory.

A possible explanation for both the reduced propulsive force and shorter angular trajectory of *disA-1* cilia is greater rotational compliance of the basal body within the power stroke plane due to the lack of SF support (Junker et al., 2022). The estimated average bending moment at *disA-1* BBs during the ciliary power stroke is reduced by 33% compared to WT (Table 1; WT: 2.86±0.87 pN·µm; *disA-1*: 1.93±1.44 pN·µm; P < 0.0001). If the BB has rotational compliance, the moment due to viscous forces on the cilium will tend to rotate it in the opposite direction of the cilium during the power stroke. We expect that the BB rotation predicted for WT cells in Fig. 1 would increase in *disA-1* cells where BBs are no longer connected. BB rotation will have the effect of reducing the swept angle for a given ciliary deformation cycle. Reducing the sweep angle alone would reduce the sweep rate for a given ciliary deformation thereby reducing the applied forces. This complex interaction requires further experimental and modeling work beyond the scope of this study. Recovery stroke parameters are largely unaffected in *disA-1* cells, perhaps because the recovery stroke occurs in a plane orthogonal to the BB axis and structural connections other than the SF are sufficient to stabilize it against moments about its axis.

### BB connections promote coordinated ciliary beating

For cilia to beat in coordination, each cilium undulates with a constant temporal delay or phase difference relative to its neighbors. BB disconnections in *disA-1* cells lead to slower, inconsistent power strokes and altered ciliary power stroke waveforms (Figs. 5A - C, S3D and I). We hypothesized that inconsistent power stroke speed and altered ciliary power stroke waveforms would disrupt the synchronization of a cilium undulating with its neighbors. To test this hypothesis, the phase difference between adjacent and oriented cilia was quantified when cilia were imaged from above the cell. The axis of the power stroke served as a proxy for BB orientation (Fig. S3A). The phase difference between cilia is influenced by the strength of their hydrodynamic coupling (Elgeti and Gompper, 2013). When cilia are positioned far apart, they are poorly hydrodynamically coupled and this leads to variable phase differences (Brumley et al., 2014). To rule out this possibility, only WT and *disA-1* cilia that were spaced apart by comparable distances were analyzed (inter-cilia spacing; WT: 2.3±1.3 µm; *disA-1*: 2.0±0.7 µm; Mann-Whitney test; P = 0.40). *disA-1* cilia exhibit more variable phase differences compared to WT cilia (Fig. S3H; phase difference; WT: −4.9±5.4 msec; *disA-1*: −0.8±14.0 msec; F-test for variance comparison; P < 0.001). Thus, BB disconnections in *disA-1* cells appear to disrupt consistent phase difference.

We next hypothesized that the inconsistent phase differences in *disA-1* mutants, would increase the probability of ciliary tangles in which neighboring cilia physically interact. Consistent with this, an 8-fold increase in ciliary tangles was observed in oriented cilia of *disA-1* cells compared to WT cells (Fig. 5K; WT: ∼20 events/ min; *disA-1*: ∼170 events/ min; Mann-Whitney test; P = 0.004). Ciliary tangles occurred when a *disA-1* cilium beat slower than its anterior and posterior neighbors during the power stroke (Movie 4 (middle and right); black arrows mark examples of this event in two *disA-1* cells). This uncoordinated behavior caused neighboring cilia within the same ciliary row to physically interfere with each other. Subsequently, these uncoordinated cilia also clash with cilia from adjacent ciliary rows that are undergoing the recovery stroke, leading to ciliary tangles between neighboring ciliary rows (Movie 4). Therefore, consistent with the biflagellate *Chlamydomonas*, BB connections are required for coordinated ciliary beating in the multi-ciliate *Tetrahymena thermophila* (Quaranta et al., 2015; Wan and Goldstein, 2016). Notably, our study suggests that BB connections promote consistent power stroke speed and waveform to minimize ciliary tangles during coordinated ciliary beating (Fig. 5L).

### Summary

Together with hydrodynamic coupling between cilia, the underlying cortical architecture of multi-ciliary arrays promotes ciliary coordination. Here we show that intracellular connections between BBs spans the cell cortex (Fig. 1). We propose that the interconnected nature of the cortical architecture transmits ciliary forces that serve as coordination cues for metachronal ciliary beating. A *Tetrahymena* cell immobilization technique was developed to show that BB connections regulate ciliary dynamics, independent of disoriented ciliary beating. BB connections promote long ciliary trajectories of cilia during the power stroke for effective fluid propulsion (Figs. 3 and 5). BB connections also promote consistent ciliary speed and waveform to maintain productive phase differences between neighboring cilia (Figs. 4 and 5). This minimizes ciliary tangles, ensuring that cilia beat in a coordinated fashion to promote efficient fluid flow and cell motility (Fig. 5L).

## SUPPLEMENTAL FIGURE LEGENDS

**Supplemental figure 1.**
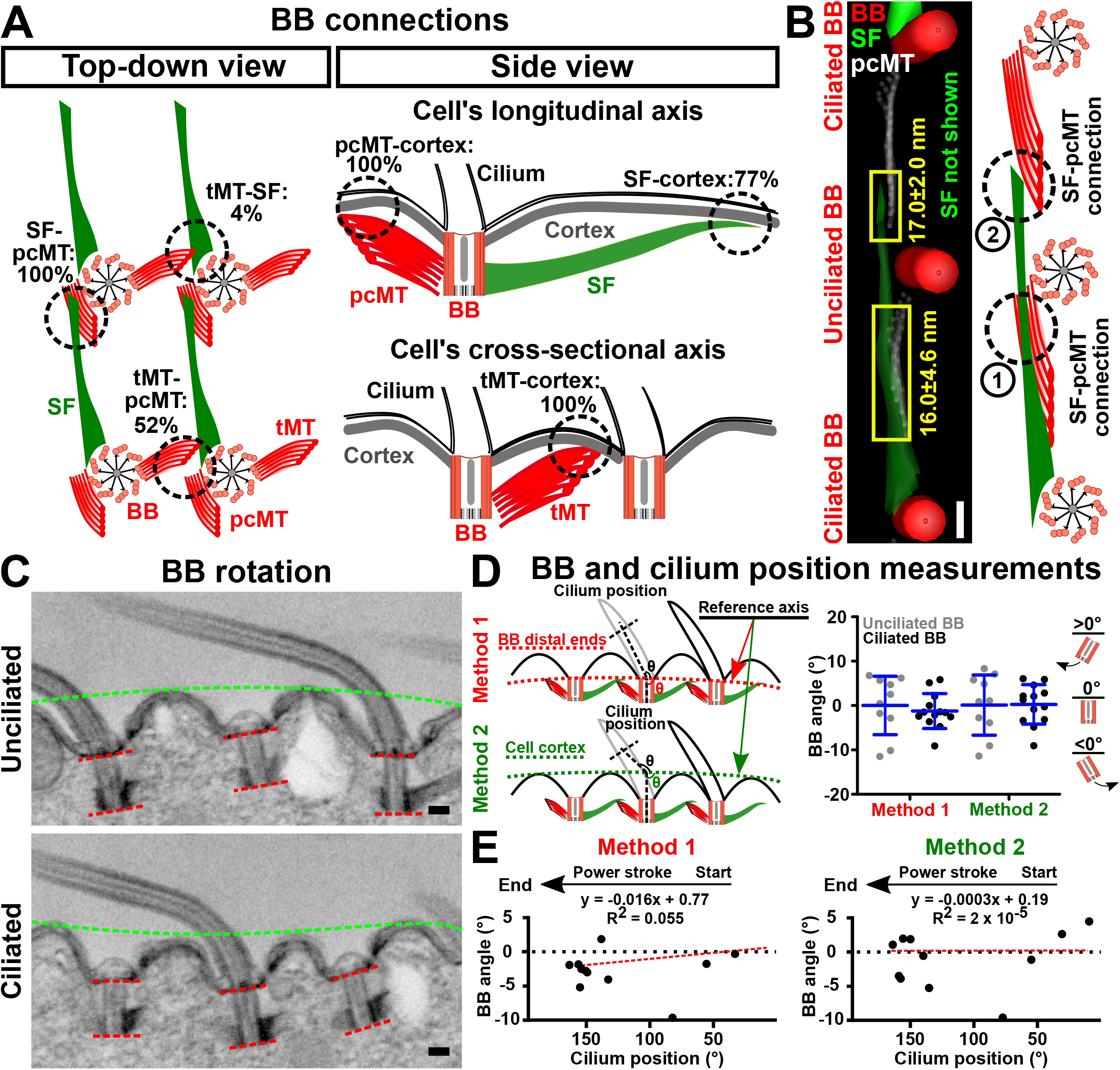
Cilia are intracellularly connected by BBs and BBs are positioned at variable angles along the cell cortex. (A) WT BB connections at 30°C. BB connections were quantified from FIB-SEM volume projections. Percentages indicate the observed frequency of BB connections. SF (n = 26). pcMTs (n = 25). tMTs (n = 27). (B) Long SFs of ciliated BBs (bottom BB) may interact with the pcMTs of both the unciliated BB (middle BB; SF not shown) and the anterior BB that is two BB units away (top BB). BB, red. SF, green. pcMT, white. Distances between SF and pcMTs at positions 1 and 2 are indicated in yellow. n = 3 BBs. Scale bar, 200 nm. (C) EM image depicting unciliated BB (top) and ciliated BB (bottom) positioning at the cell cortex. Cell cortex (green dashed line) and BB distal and proximal ends (red dashed lines) are marked to reflect BB positioning relative to the cell cortex. Scale bars, 200 nm. (D) *Left panel*: Schematic illustrates the analysis methods to quantify BB positioning along the cell cortex. Reference axes across the distal end of three consecutive BBs (method 1) and across peaks along the cell cortex (method 2) were used. *Right panel*: Quantification of BB angle using both methods. BBs are docked at variable angles along the cell cortex. n = 5 cells. Unciliated BBs: 10. Ciliated BBs: 13. (E) Correlation between cilium position along the power stroke axis and the BB position along the cell cortex quantified using analysis methods 1 and 2. Cilium position along the power stroke axis is defined by measuring the angle from 1.5 µm − 2.0 µm up the cilium relative to the reference axes.

**Supplemental figure 2.**
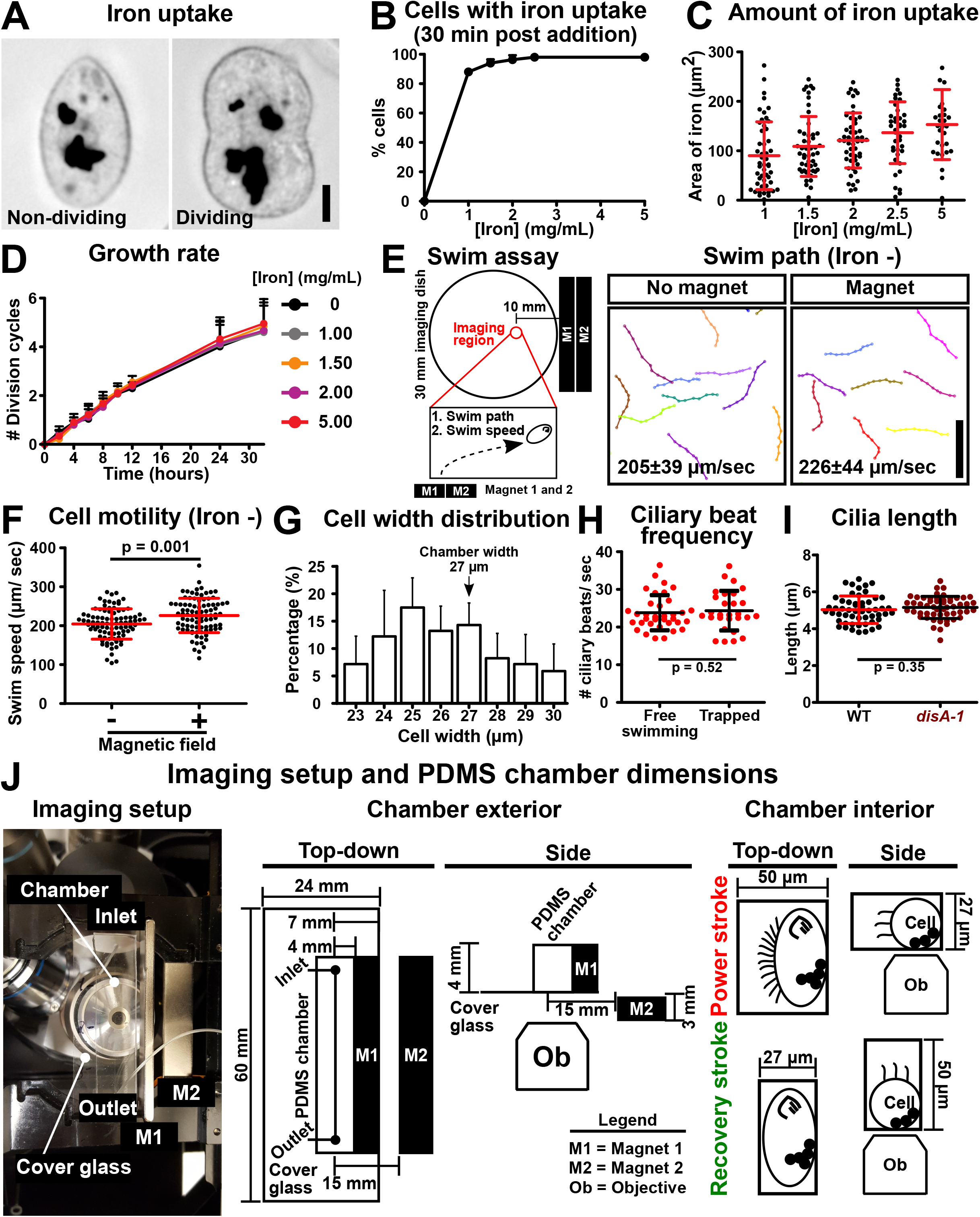
Immobilization of live *Tetrahymena* cells using DIPULL microscopy. (A) Phagocytosis of iron particles by non-dividing and dividing cells. Scale bar, 10 µm. (B) Percentage of cells with iron uptake after exposure to different concentrations of iron particles. Quantitation was performed at 30 min post iron feeding. n = 150 - 250 cells. (C) Area (2-dimensional) of phagocytosed iron particles when cells were fed with varying concentrations of iron particles for 30 min. n = 30 - 50 cells. (D) Growth rate of *Tetrahymena* cells is not inhibited by the range of iron concentrations tested. (E and F) In the presence of the applied magnetic field used in DIPULL microscopy, *Tetrahymena* cells’ swim trajectory is not affected but they swim slightly faster. Student’s t-test. n = 90 cells. (G) Quantification of *Tetrahymena* cell width to determine microfluidic chamber width for long-term live cell imaging. n = 179 cells. Chamber width of 27 µm was determined to accommodate most cells. (H) DIPULL does not inhibit *Tetrahymena* ciliary beat frequency. Free-swimming n = 23 cells. Immobilized n = 47 cells. Mann-Whitney test. Mean±SD. (I) Quantification of WT and *disA-1* cilia length. n = 18 cells (3 cilia each). Student’s t-test. Mean±SD. (J) Imaging setup and microfluidic chamber design.

**Supplemental figure 3.**
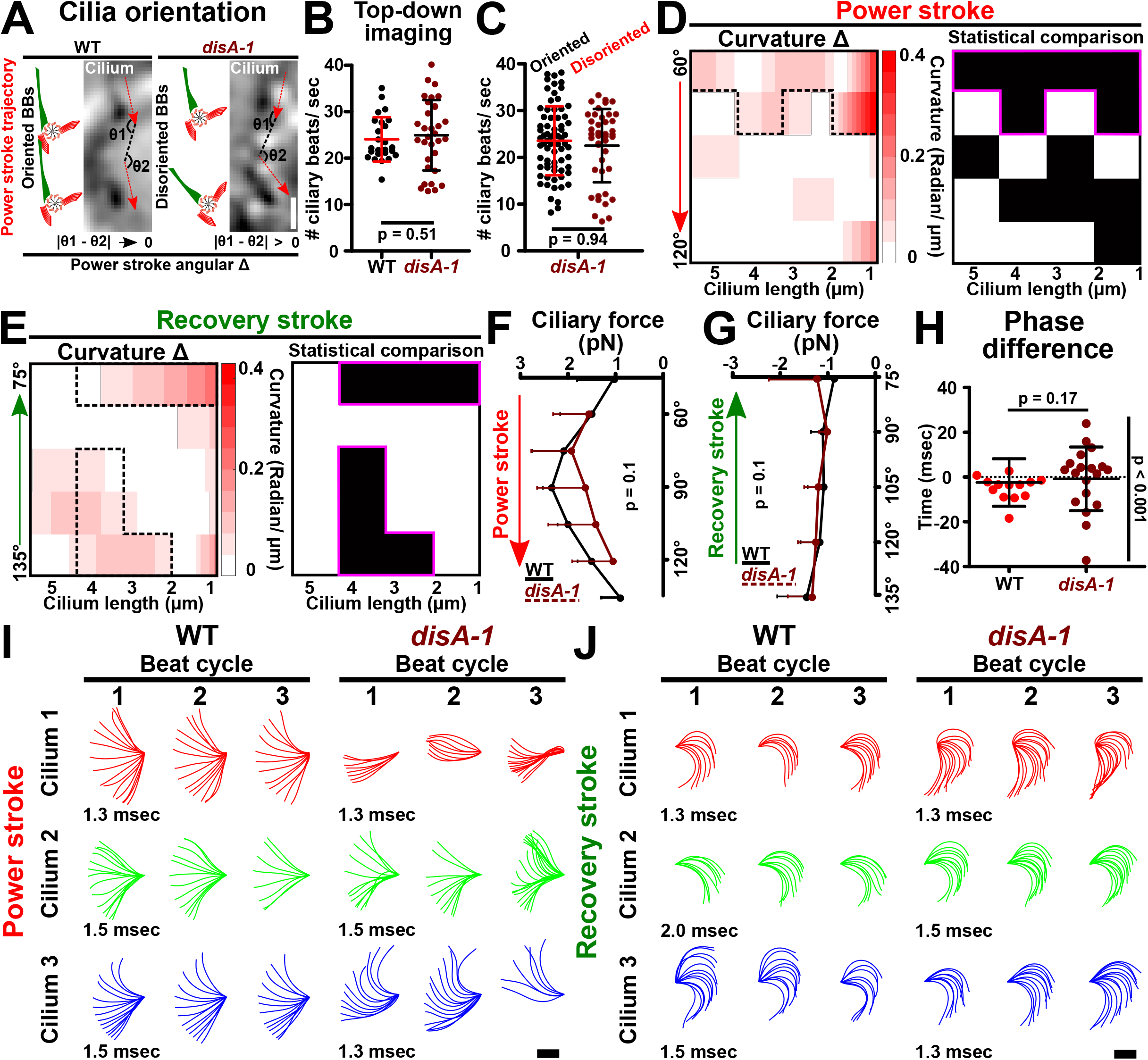
BB connections support ciliary waveform and coordination. (A) Schematic illustrates the power stroke trajectories of two adjacent cilia imaged from the top of the cell. BB orientation is determined based on the cilium’s power stroke axis between adjacent cilia. If adjacent cilia are oriented, the power stroke angular difference is close to zero. Conversely, disoriented adjacent cilia would have large power stroke angular differences. Bar: 1 µm. (B) Quantification of ciliary beat frequency using cilia that were imaged from the top of the cell. *disA-1* ciliary beat frequency is comparable to WT cilia. WT: n = 11 cells (25 cilia). *disA-1*: n = 13 cells (32 cilia). Mann-Whitney test. Mean±SD. (C) Oriented and disoriented cilia display comparable ciliary beat frequencies. *disA-1*: n = 38 cells (72 oriented cilia; 41 disoriented cilia). Mann-Whitney test. Mean±SD. (D) Net curvature difference and statistical comparison between WT and *disA-1* cilia during the power stroke. *disA-1* cilia are more bent in the medial and proximal region between cilium position 60° - 75° (black dotted outline and pink outline). Absolute values of curvature difference are indicated (white: zero curvature difference; red: large curvature difference). Black indicates regions of the cilium where the curvature difference is statistically significant. Student’s t-test. P value < 0.01. (E) Net curvature difference and statistical comparison between WT and *disA-1* cilia during the recovery stroke. *disA-1* cilia are slightly more bent at cilium positions 75°, and 120-135° (black dotted outline and pink outline). Absolute values of curvature difference are indicated (white: zero curvature difference; red: large curvature difference). Black indicates regions of the cilium where the curvature difference is statistically significant. Student’s t-test. P value < 0.01. (F) Ciliary power stroke forces were estimated using resistive force theory. WT cilia exert more power stroke force per stroke than *disA-1* cilia (n = 9 cilia; 6 ciliary beat cycles). Data bin size is 15°. (G) Ciliary power stroke forces were estimated using resistive force theory. WT cilia exert comparable recovery stroke force per stroke relative to *disA-1* cilia (n = 9 cilia; 6 ciliary beat cycles). Data bin size is 15°. (H) Quantification of WT and *disA-1* phase differences. WT: n = 11 cells (13 cilia pairs). *disA-1*: n = 13 cells (19 cilia pairs). Mann-Whitney test; F-test for variance comparison. Mean±SD (I) Individual power stroke waveforms of WT and *disA-1* cilia. Representative examples of WT and *disA-1* power stroke waveforms showing three consecutive beat cycles. *disA-1* cilia display variable power stroke waveforms. Imaging frame intervals are indicated (msec). Scale bar, 2 µm. (J) Individual recovery stroke waveforms of WT and *disA-1* cilia. Representative examples of WT and *disA-1* recovery stroke waveforms showing three consecutive beat cycles. Imaging frame intervals are indicated (msec). Scale bar, 2 µm.

## MOVIE LEGENDS

**Movie 1. FIB-SEM image volume illustrating WT *Tetrahymena* cortical architecture.** Cilia, BB, MT appendages, red. SF, green. Scale bar, 1 µm.

**Movie 2. WT and *disA-1* fluid flow dynamics.** Imaging frame interval, 40 fps. Movie frame rate, 20 fps. Scale bar, 50 µm.

**Movie 3. WT and *disA-1* ciliary power stroke.** Cell anterior pole is oriented to the top of the movie. Disoriented cilia do not beat in a uniform direction, and they beat out of the imaging plane. Imaging frame interval, 793 fps. Movie frame rate, 20 fps. Scale bar, 5 µm.

**Movie 4. WT and *disA-1* ciliary recovery stroke.** Cell anterior pole is oriented to the top of the movie. Ciliary tangles within and between ciliary rows occur when *disA-1* cilia (black arrow) undergo inconsistent power stroke. Examples of ciliary tangles in two *disA-1* cells are provided. Imaging frame interval, 793 fps. Movie frame rate, 20 fps. Scale bar, 5 µm.

## Materials and methods

### *Tetrahymena* culture

*Tetrahymena thermophila* cells were grown in 2% SPP media (2% proteose peptone, 0.2% glucose, 0.1% yeast extract, and 0.003% Fe-EDTA). All studies were performed under cycling conditions. Cells were grown to mid-log phase (2.5 - 4 x 10^5^ cells/ ml) prior to immobilization. Cell concentration was determined using a Coulter Counter Z1 (Beckman Coulter). All analyses were restricted to non-dividing cells as judged by the absence of a developing oral structure.

### *Tetrahymena* immobilization (DIPULL-microscopy)

*Tetrahymena* cells are fed iron particles (Sigma-Aldrich; 267953-5G (particle size: < 10 µm); Figs. 2A and S2A; (Rifkin and Ballentine, 1976)). To determine the ideal iron concentration for DIPULL, a growth assay was performed on cells that were fed with a range of iron concentrations (1 - 5 mg/ ml). *Tetrahymena* cells readily phagocytose iron particles and the amount of engulfed iron is concentration-dependent (Fig. S2B and C). The tested iron concentrations applied in DIPULL do not disrupt *Tetrahymena* cell growth or morphology (Fig. S2A and D). For consistency, cells were supplemented with an iron concentration of 2.5 mg/ ml (suspended in 10 mM TRIS-HCL, pH 7.4) for all live cell imaging experiments performed in this study. Final media concentration was 1.3% SPP and 3.3 mM TRIS-HCL (pH 7.4).

Next, *Tetrahymena* cells containing phagocytosed iron were introduced into microfluidic chambers of specified dimensions (Fig. 2A and S2J). For extended live cell imaging, microfluidic chambers with channels that constrain *Tetrahymena* cells were used (Fig. S2G and J; Minimum chamber width and depth: 27.0 and 27.0 µm). To study ciliary dynamics, two different microfluidic chambers were used. To image the power stroke plane, iron particle-fed *Tetrahymena* cells were introduced into wide chambers with narrow depth (Fig. S2J; Chamber dimensions: 50.0 µm x 27.0 µm; cell width: 26.0 ± 2.7 µm; cilia length: 5.0 ± 0.7 µm; Mean±SD). The wider axis of the chamber allows for unimpeded ciliary beating along the side of the cell. The shallow depth restricts cellular motion in the axial axis and maintains the cell within the same imaging plane. To image the recovery stroke plane, a deeper chamber was utilized to image cilia positioned at the top of the cell to avoid boundary effects (away from the cover glass) (Fig. S2J; Chamber dimensions: 27.0 µm x 50 µm; cell width: 26.0 ± 2.7 µm; cilia length: 5.0 ± 0.7 µm; Mean±SD).

To immobilize *Tetrahymena* cells, a constant external magnetic field using two bar magnets was applied (X-bet MAGNET N52 grade; Fig. S2J). Magnetic fields were previously shown to disrupt *Paramecium* motility (Nakaoka et al., 2002). To establish optimal magnetic field strength for DIPULL, we investigated *Tetrahymena* swim path in the presence of a constant magnetic field exerted by the two bar magnets. *Tetrahymena* cells display normal swim paths and a slightly elevated swim speed in the presence of the applied magnetic field (Fig. S2E and F). When cells were not stably immobilized via two magnets, an additional magnet was placed below the microscope stage insert to further stabilize *Tetrahymena* immobilization without impacting ciliary beat frequency (Fig. S2H). By coupling magnetism and microfluidic chambers to achieve cell immobilizations, DIPULL microscopy is applicable for short- or long-term live cell imaging.

### Microfluidic chamber fabrication

Microfluidic immobilization chambers were fabricated in polydimethylsiloxane (PDMS), which has been shown to be salubrious for hosting long-term cell culture and imaging (Bascom et al., 2016). Standard soft lithography protocols (Xia and Whitesides, 1998) were used to produce PDMS chambers, which were bonded to cover glass to enable imaging by light microscopy. Briefly, negative photoresist (SU 8, Kayaku Advanced Materials, Inc., Westborough, MA) was spun upon a silicon wafer to the desired depth, baked, and exposed to channel patterns by illuminating with collimated UV light through a shadow mask. Rinsing and washing in developer solution revealed a positive mold of the channel network, which was then covered with uncured, degassed PDMS and baked for 70°C for four hours to cure. Devices were completed and bonded by trimming the *bas* relief replica, punching holes for fluid and cell introduction and removal, exposing to an oxygen plasma (Herrick), and placing in contact with a clean cover slip.

### Light microscopy

Imaging experiments were performed with an inverted widefield microscope (Ti Eclipse; Nikon). Depending on the experiment, a 20x objective lens (Ti Eclipse; Nikon) or 60xA DIC Plan-Apo (NA 1.40) objective lens (Nikon) was used. Images were captured with a scientific complementary metal-oxide semiconductor (CMOS) camera (Zyla; Andor Technology).

### I. Visualizing Tetrahymena ciliary mobility

Due to the 3-dimensional waveform of *Tetrahymena* cilia, two DIC imaging regimes were used to track the power and recovery strokes along the XY-axes of the imaging plane. To visualize the power stroke, cilia mobility along the length of the cell was imaged (Fig. 2B and Movie 3). To track the recovery stroke, cilia positioned on the top of the cell were imaged (Fig. 2B and Movie 4). Imaging frame rates were between 1.3 - 2.0 msec.

### II. Imaging fluid flow dynamics

To track fluid flow, the displacement of 0.5 µm fluorescent beads in the SPP growth media (Polysciences Inc; 18339) was imaged using a 20x objective lens (Ti Eclipse; Nikon). To improve signal:noise, movies were acquired with 3×3 binning. Each movie duration was 12.5 seconds (imaging frame rate: 40 frames per second).

### Focused Ion Beam Scanning Electron Microscopy (FIB-SEM)

High pressure freezing and freeze substitution (HPF-FS) and sample preparation were prepared as previously described (Giddings et al., 2010; Meehl et al., 2009; Soh et al., 2020). Heavy metal stained, resin-embedded samples were sectioned, and consecutive sections were screened by TEM imaging to identify cells appropriate for FIB-SEM (embedded cells with intact morphology and that were not sectioned). The specimen was trimmed, mounted on a stub, and introduced into a Zeiss Crossbeam 550 FIB-SEM instrument. Image acquisition and processing followed a general schema outlined in our prior study (Baena et al., 2021). Briefly, SEM imaging of the sectioned face allowed the re-localization of these cells, which were protected with a patterned platinum and carbon pad and exposed orthogonally by controlled FIB milling up until the appearance of the tips of cilia. Automated FIB-SEM acquisition cycles of milling and imaging were executed over a period of several days, with SEM image pixel sampling of 5 nm in the imaging plane and 15 nm FIB mill “z step” size. The resulting stack of hundreds of images was aligned, contrast inverted and binned by a factor of 3 to yield 8 bit, isotropic 15 nm voxel image volume reconstructions. Sub-volumes of interest from these were then segmented and analyzed.

### *Tetrahymena* cell image analysis

#### I. Quantification of BB angle at the cell cortex

Undulation at the cell cortex may cause erroneous interpretations of BB docking orientation (Figs. 1B and S1C). To rule out non-uniformity along the cell cortex, two reference axes were utilized for BB angle measurements. In method 1, a reference axis that spans across the distal ends of two flanking BBs and the BB of interest was utilized (Fig. S1D; straight line was fitted through the midpoint on the distal end of three consecutive BBs). In method 2, the highest points of the cell cortex that span across three consecutive BBs were used as an alternate reference axis (Fig. S1D; straight line was fitted through the highest points along the cell cortex). The BB angle relative to the cell cortex was determined from the angle between the BB longitudinal axis and the two reference axes (Methods 1 and 2). To establish whether the BB angle depends on the cilium’s position, the position of the cilium along the power stroke was quantified by measuring the angle from 1.5 - 2.0 µm up the cilium relative to the reference axes.

#### II. Quantification of Tetrahymena iron phagocytosis

Fluorescence quantification of iron uptake was performed by a semi-automated strategy that utilizes the FIJI macro scripting language and plugins (Schindelin *et al*., 2012). Images (2-dimensional) were preprocessed by a local background subtraction. To obtain the area of the engulfed iron particles, a uniform intensity threshold was applied to generate a binary mask over iron particles. The area of the binary mask served as a proxy for the area of ingested iron particles.

#### III. Tetrahymena motility analysis

*Tetrahymena* cell motility was imaged on an inverted widefield microscope using a 20x objective lens (Ti Eclipse; Nikon). Each movie is 2.5 sec in duration (frame rate: 20 frames per second). To quantify swim rates, the relative displacement of the anterior or posterior pole of cells was tracked for 500 msec via the FIJI MTrackJ plugin (Meijering *et al*., 2012). Analyses were restricted to cells that swim along the same XY plane.

#### IV. Cilia length measurement

To measure cilium length, the FIJI freehand tool was used to trace from the ciliary base to tip. Live cilia movies acquired by DIC microscopy were used in this analysis. Analysis was restricted to cilia positioned at the cell’s medial region. Six cells (three cilia per cell) were analyzed.

#### V. Ciliary kinetics measurements

To quantify cilia beat frequency, kymograph analysis on movies of *Tetrahymena* ciliary beating acquired by DIC microscopy was performed. Kymograph analysis was performed on cilia that were positioned along the cell’s medial region (Fig. 2B; red box). The number of in-focused cilia at the cell’s medial region is usually three. To ensure equal weightage, three cilia per cell were analyzed. For each cilium, three consecutive beat cycles were followed. To assess BB orientation along the side of the cell, three criteria were imposed on the cilia that were selected for this analysis. First, cilia must transverse along the cell’s anterior-posterior axis. Second, cilia must remain in-focus during the entire power stroke before transitioning into the recovery stroke where they exit the imaging focal place (Fig. 2B; right). Third, the anteriorly and posteriorly positioned neighboring cilia must also remain in-focus during analysis (Movie 3). Kymographs were generated from an 11-pixel (pixel size: 108 nm) wide line that was positioned approximately 1.5 µm from the ciliary base. Using FIJI, three kymographs (one per cilium) were generated for each cell (FIJI Multi Kymograph plugin). Each ciliary beat cycle is represented as a peak on the kymograph whereby the upward slope marks the power stroke while the downward slope marks the recovery stroke (Fig. 4A). Based on the duration of three consecutive beat cycles, the average cilia beat frequency (per second) was quantified.

To provide better assessment of BB orientation, kymograph analysis was also performed on cilia that were imaged from the top of the cell (Fig. 2B; green box). To quantify BB orientation, the difference in angle along the power stroke axis between two adjacent cilia, or power stroke angular difference, was measured (Fig. S3A). An average power stroke angular difference was quantified from each cilia pair across three consecutive ciliary beat cycles. WT cilia pairs exhibit a power stroke with an average angular difference of 3.7±16.0°. *disA-1* cilia pairs were considered oriented if their power stroke angular differences fall within 1 SD of the average WT power stroke angular difference. To avoid boundary effects from the imaging chamber, cilia that were positioned away from the chamber ceiling were quantified. Ciliary beat frequencies are comparable between cilia that were imaged from the side view and top-down view.

To quantify the duration of the power and recovery strokes, kymograph analyses were performed on cilia that were imaged from the top of the cell. Average power and recovery stroke duration was calculated from three consecutive ciliary beat cycles. The phase difference between adjacently positioned cilia was also quantified using kymograph analyses. To measure the temporal delay between when a cilium undergoes its ciliary beat cycle relative to its neighbor, kymographs were generated from an 11-pixel (pixel size: 108 nm) wide line that bisected two adjacent and in-focused cilia. In addition, the line was positioned at approximately 1.5 µm from the ciliary base of both cilia. An average phase difference was calculated from three consecutive ciliary beat cycles.

#### VI. Ciliary force estimation

##### Calculation of forces based on ciliary waveform analysis using resistive force theory

Forces on *Tetrahymena* cilia were calculated using resistive force theory (Gray and Hancock, 1955). Estimates for the value of the resistive force coefficient vary from approximately 1 - 3.5-fold the viscosity of the fluid (Bayly et al., 2011; Lighthill, 1976; Riedel-Kruse et al., 2007). We used the values obtained for wild-type *Chlamydomonas reinhardtii* cilia as previously described (Bayly et al., 2011); *C_N_=1.53×10^-3^pN·s/µm^2^*, *C_T_=0.64×10^-3^pN·s/µm^2^*)). These values were corrected to account for changes in the viscosity of the surrounding fluid due to temperature differences.

The average ciliary waveform was generated using a previously published protocol (Bottier et al., 2019). To ensure equal weightage, each cilium was followed for six consecutive beat cycles. Nine cilia (each from a different cell) were analyzed. Briefly, consecutive frames of *Tetrahymena* cilia undergoing a power stroke (XY) and a recovery stroke (YZ) were manually traced (Fig. 2B). Due to poor image contrast at the cilium proximal end (∼1.5 µm), this study was unable to accurately determine the position of the cilium base. Images were processed to improve cilia contrast using FIJI Unsharp mask (radius: 1 pixel; mask weight: 0.9). For the power stroke, each cilium was aligned along the cell’s anterior-posterior axis using the base of the anterior cilium as a reference point. For the recovery stroke, each cilium was aligned relative to the power stroke axis of the same cilium and the cell’s anterior-posterior axis. Next, each trace was fitted with a polynomial function. Two-dimensional average ciliary waveforms of the power stroke and recovery stroke were generated by averaging across a large number of individual ciliary traces (6 beat cycles of 9 cilia total). The cilium position is defined by its orientation from the cilium’s distal end (4.5 µm up from the cilium base) relative to the cilium’s base and cell’s anterior pole. The power stroke was divided into three stages based on the orientation of the cilium relative to the cell’s anterior-posterior (AP) axis (start: 0-60°; middle: 61-120°; end: 121-180°). The recovery stroke was divided into three stages based on the orientation of the cilium relative to the cell’s posterior pole (start: 180 - 121°; middle: 120 - 61°; end: 60 - 0°). Cilium positions are categorized into 15° bins for both the power and recovery strokes. The start and end positions of each cilium along the power stroke and recovery stroke were defined by the orientation of the first and last ciliary trace that appear along the imaging focal plane. The reference point for angle measurement is 4.5 µm up from the cilium base relative to the cell’s anterior-posterior axis. The average start and stop positions for each cilium were calculated by averaging the start and end positions across 6 ciliary beat cycles. The angular trajectory for both the power and recovery strokes were quantified by measuring the angle between the average start and end positions of each cilium. Analyses were focused on ciliary traces that fell within the average angular trajectories (Fig. 5B and G). The net difference in ciliary curvature between WT and *disA-1* cilia was computed by subtracting the average curvature of *disA-1* from that of WT. Absolute values of the net difference in ciliary curvature is presented (Fig. S3D and E).

Cilium velocity, *v_c_*(*s*,*t*), as a function of arclength coordinate, *s*, and time, *t*, was calculated by numerically differentiating the coordinates of the cilium traces with respect to Velocities are projected onto the tangent and normal vectors along the cilium for calculation of the distributed resistive forces (Bayly et al., 2011). Power per unit length (fW/μm) along the cilium is calculated as the dot product of the distributed force and velocity, *P*(*s*,*t*) = *F*(*s*,*t*) · *v*(*s*,*t*).

Total power produced by the cilium at a given time is obtained by numerically integrating the power over the length of the cilium. Moment at the cilium basal attachment (pN · μm) is calculated as the integral along the length of the cilium of the cross product of the position relative to the basal attachment and the distributed force, 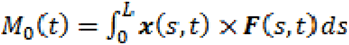.

### Calculation of forces on the cell body

Forces on the cell body were calculated for comparison against ciliary forces estimated from RFT. Average swimming velocities for WT and *disA-1* cells used in calculations were based on previously reported study (Galati et al., 2014). The cell body was modeled as an ellipsoid moving along its major axis (Bayly et al., 2011; Chwang and Wu, 1975). To estimate the average force each cilium, the body drag force along the cell’s anterior-posterior axis was divided by the estimated number of cilia, 400 (assuming all BBs are ciliated; (Galati et al., 2015)).

#### I. Fluid flow analysis

To quantify fluid flow dynamics, we measured the displacement of fluorescent beads by immobilized *Tetrahymena* cells using PIVlab (Thielicke, 2021). Fifty consecutive image frames were first processed using the wiener2 denoise filter (3 pixel; imaging frame rate: 40 frames per second). Depending on the signal:noise of the movies, two interrogation windows (64 pixel only or both 64 and 32 pixels) were utilized to track fluorescent bead motility. An averaged vector map that depicts fluid flow velocity was generated for each cell. To measure fluid flow along the length of the cell, the polyline tool was used to measure fluid velocity along the anterior-posterior axis of the cell. Fluid flow was visualized using FIJI Flowtrace plugin (Gilpin et al., 2017a).

### Statistical analysis

All datasets were assessed for normal distribution using the Shapiro-Wilk or Kolmogorov-Smirnov test. A student’s t-test was performed on normally distributed datasets. A Mann-Whitney test was performed on datasets that do not conform to a normal distribution. An F-test was performed to compare variance between conditions. Tests for significance were unpaired and two-tailed. All error bars indicate SD. P value is indicated for all statistical analyses. All analyses were performed on samples obtained from 3 independent experiments.

## Acknowledgements

We thank the Pearson lab for the helpful and enjoyable discussions during this project. We also express our gratitude to Alexander J. Stemm-Wolf for critical reading of the manuscript. We are thankful to Dr. Tom H. Giddings, Eileen T. O’Toole, and Garry Morgan (University of Colorado Boulder) for EM sample preparation and sample screening used in FIB-SEM imaging. Research was funded by the National Institutes of Health-National Institute of General Medical Sciences (R01GM099820 and R35GM140813), the Pew Charitable Biomedical Scholars Program, and the W.M. Keck Foundation (C.G. Pearson). C.M.E. acknowledges a graduate fellowship from the NIH-funded Wyoming IDeA Networks of Biomedical Research Excellence program (P20GM103432). This project is also funded in part with Federal funds from the National Cancer Institute, National Institutes of Health, under Contract No. 75N91019D00024. The content of this publication does not necessarily reflect the views or policies of the Department of Health and Human Services, nor does mention of trade names, commercial products, or organizations imply endorsement by the U.S. Government.

